# Structure of the Flotillin Complex in a Native Membrane Environment

**DOI:** 10.1101/2024.05.09.593390

**Authors:** Ziao Fu, Roderick MacKinnon

## Abstract

In this study we used cryo-electron microscopy to determine the structures of the Flotillin protein complex, part of the Stomatin, Prohibitin, Flotillin, and HflK/C (SPFH) superfamily, from cell-derived vesicles without detergents. It forms a right-handed helical barrel consisting of 22 pairs of Flotillin1 and Flotillin2 subunits, with a diameter of 32 nm its wider end and 19 nm at its narrower end. Oligomerization is stabilized by the C-terminus, which forms two helical layers linked by a β-strand, and coiled-coil domains that enable strong charge-charge inter-subunit interactions. Flotillin interacts with membranes at both ends; through its SPFH1 domains at the wide end and the C-terminus at the narrow end, facilitated by hydrophobic interactions and lipidation. The inward tilting of the SPFH domain, likely triggered by phosphorylation, suggests its role in membrane curvature induction, which could be connected to its proposed role in clathrin-independent endocytosis. The structure suggests a shared architecture across the family of SPFH proteins and will promote further research into Flotillin’s roles in cell biology.

**Significance statement:** It is well known that many biochemical processes in cells must occur in localized regions. There are many different ideas about how cells keep processes localized. In this study we demonstrate that Flotillin1 and Flotillin2 co-assemble to form a large, truncated cone shaped cage whose wide end is always attached to a membrane surface and whose narrow end is sometimes attached to a separate membrane. The entire wall of the cage is without holes and is likely impervious even to small molecules, forming a diffusion barrier that can connect membrane systems. The Flotillin cage is thus well suited to isolate biochemical processes. Through membrane attachment, it also alters local membrane curvature, which could influence endocytic and mechanosensory processes.

## Introduction

While analyzing plasma membrane vesicles isolated from HEK293 GnTl^−^ cells using cryo-electron microscopy we observed large, truncated cone shaped objects attached to membrane surfaces and wondered what they were. We looked in other cell types, including red blood cells, and observed similar objects. We determined the molecular structure of these objects and report it here. For reasons that will become clear, we call these structures Flotillin cages.

Flotillin1 and Flotillin2, also called reggie2 and reggie1^1,2^, respectively, are mammalian members of a protein family called SPFH proteins, named for Stomatin, Prohibitin, Flotillin and HlfK/C, which all contain namesake SPFH domains^3^. The name Flotillin originates in their discovery among proteins floating near the top of density gradients during the fractionation of detergent cell lysates^1^. Flotillins have been implicated in many different cellular processes in the cell biological and biochemical literature, including caveolae function^1^, endocytosis independent of clathrin and caveolae^4–6^, IgE receptor mediated signaling^7^, and cholesterol uptake and trafficking^8^. A common theme that emerges from the rather extensive literature on these proteins is that they somehow mediate the formation of membrane microdomains^8–13^

In 2022 a first structure of an SPFH family protein, a complex of HflK and HflC from bacteria was determined^14^. Twenty-four subunits, 12 HflK and 12 HflC in an alternating arrangement, form a cage with a wide and narrow end. The wide end was attached to a detergent micelle by 24 transmembrane helices, one from each subunit. The cage contained inside it, at the wide end facing the micelle, between 0 and 4 hexameric AAA+ proteases. The authors proposed that the HflK/HflC complex docks onto the membrane and forms a new kind of microdomain that could insulate the biochemical process inside the cage (proteolysis) from the surrounding membrane. They further hypothesized that other SPFH proteins may function as membrane delimiting microdomains.

In this study we describe our findings on the structure of a second member of this protein family, a complex formed by the mammalian proteins Flotillin1 and Flotillin2.

## Results

### Flotillin Cages on Cell Derived Membranes

Figure 1A outlines three separate methods we used to prepare membrane vesicles in this study. In one method we produced giant plasma membrane vesicles by applying N-ethylmaleimide (NEM) and calcium^15^ and sonicated them to reduce their size for cryo-EM analysis^16^. In another, we isolated membrane fragments without NEM and calcium application as described in methods. In a third, we isolated spontaneous exosomes and sonicated them. All three methods yielded vesicles with Flotillin cages attached to membranes. We note that the first and third methods presumably produce vesicles originating from the plasma membrane, whereas the second method produces vesicles from both plasma and organellar membranes. As depicted in Figure 1A and shown in Figure 1B and Figure S1B-D, Flotillin cages sometimes attach to the inside of a vesicle and sometimes to the outside. We learned while studying other proteins in these kinds of vesicles that plasma membrane vesicles probably start out with an outside-out orientation, but the orientation in some vesicles can reverse with sonication.

**Fig 1.**
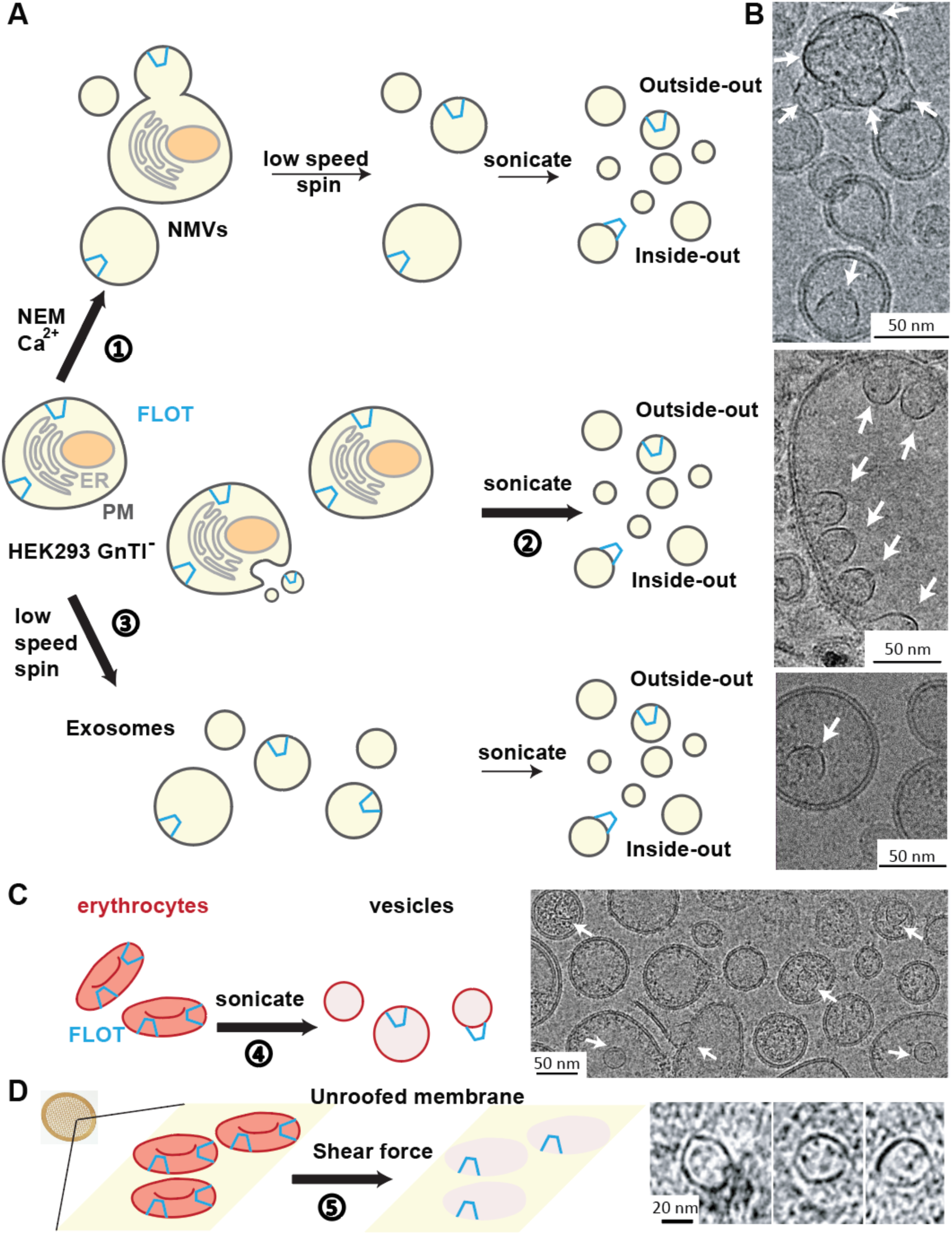
Flotillin Vesicle Preparation from Various Cell Membranes. **(A)** Schematic representation of three methods employed for the preparation of Flotillin vesicles from HEK293 GnTl-cells. The endoplasmic reticulum (ER) membrane is depicted in light gray, the plasma membrane (PM) in dark gray, the nuclei in brown, and Flotillin (FLOT) in blue. (1) Preparation of Flotillin vesicles from the plasma membrane using NEM and Ca2+, resulting in native membrane vesicles (NMVs). Following sonication, two orientations of Flotillin were observed: Inside-out, where Flotillin is outside the vesicles, and outside-out, where Flotillin is inside the vesicles. (2) Procedure for obtaining total membrane vesicles primarily through sonication. (3) Method for isolating exosomes secreted from HEK293 GnTl^−^ cells. **(B)** Representative micrographs illustrating Flotillin vesicles from the plasma membrane (top), total membrane (middle), and exosomes (bottom). Flotillin clusters were observed within vesicles, facing either outside (top) or inside (middle) the vesicle. Scale bar: 50 nm. White arrows highlight Flotillin. **(C)** Flotillin presence in erythrocyte vesicles generated by sonication, depicted in a representative micrograph (right). Erythrocytes are colored in red, Flotillin in blue. Scale bar: 50 nm. White arrows highlight Flotillin. **(D)** Observation of Flotillin in cryo-electron tomograms of erythrocyte membrane patches unroofed by shear force. Examples from the same membrane patches show erythrocytes stuck on the cryo-EM grid support (yellow surface) unroofed using shear force. The bottom membrane (pink) of erythrocytes containing Flotillin was imaged using cryo-electron tomography, with tomographic slices of Flotillin shown on the right. Scale bar: 20 nm.

The Flotillin cages occur naturally in HEK293 GnTl^−^ cells as no exogenous genes were expressed in this study. We observed Flotillin cages in other cells as well, for example A431 cells and human erythrocytes. Figure 1C shows images of erythrocyte vesicles after removal of hemoglobin and cytoskeletal elements followed by sonication. Flotillin cages inside the vesicles look like those from the HEK293 GnTl^−^ cells. Whole erythrocytes were also treated with a hypotonic buffer to induce swelling and then unroofed on a cryo-EM grid using fluid flow mediated shear force. Cryo-EM tomographic images of the erythrocyte membrane patches remaining on the grid showed membrane-attached Flotillin cages (Figure 1D).

It is likely that vesicles with exterior Flotillin cages, e.g., Figure 1A and B, are flipped inside-out vesicles and that the cages are naturally present on the cytoplasmic face of plasma membranes and possibly organellar membranes as well.

### Structure of a Flotillin Complex

We processed images from vesicles prepared using the NEM, Ca^2+^ method of GPMV formation followed by sonication (Figure 1A and Methods). From a dataset consisting of 19,024 micrographs, we manually selected Flotillin cages in numerous orientations. Top views show a circular structure with a diameter of about 30 nm. Side views resemble a truncated cone (Figure 2A-C, Fig. S1B-D). The wide end of the truncated cone is always attached to a membrane surface and typically alters the curvature of the membrane at the attachment region relative to that of the surrounding membrane. We will show later that the curvature of the membrane also alters the shape of the Flotillin cage. Thus, Flotillin and the membrane mechanically interact with each other. 40% of Flotillin cages reside inside vesicles (outside-out vesicles) and 60% reside outside vesicles (inside-out vesicles). Many Flotillin cages on the outer surface of inside-out vesicles contain a second, smaller vesicle attached to the opposite end of the truncated cone (Figure 2AB).

**Fig 2.**
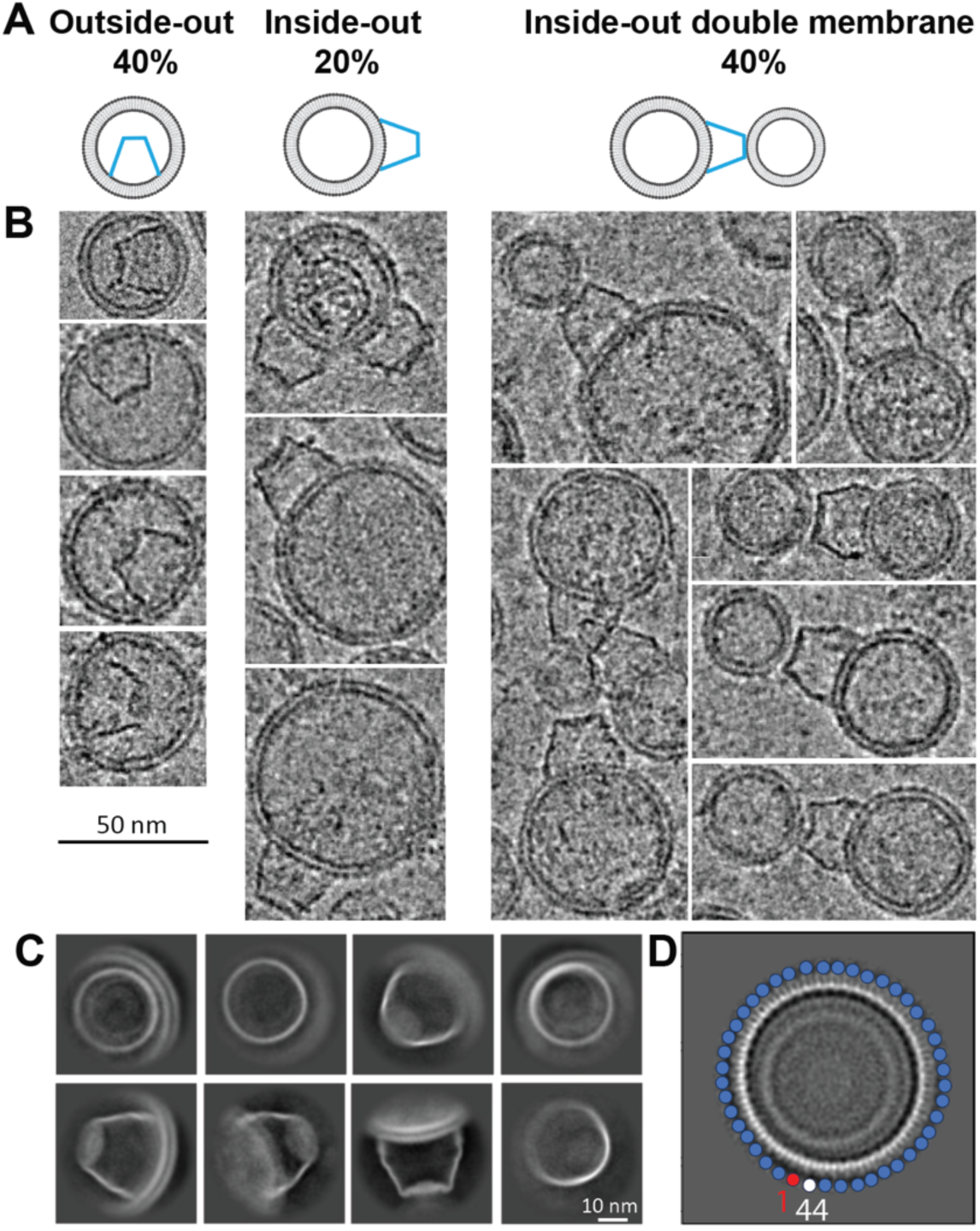
Flotillin Orientation Relative to Vesicles. **(A)** Sideview of Flotillin vesicles illustrates three cases: 40% of Flotillin is inside the vesicle (referred to as outside-out, left), while 60% of Flotillin is outside the vesicles (inside-out). In the inside-out configuration, one-third of the Flotillin have a free narrow end, and two-thirds bind to a second vesicle at the narrow end. **(B)** Examples of Flotillin vesicles showcasing outside-out (left), inside-out (middle), and inside-out double membrane (right) orientations. Scale bar: 50 nm. **(C)** Representative 2D classifications of Flotillin from vesicles displaying various orientations. Scale bar: 10 nm. **(D)** High-resolution 2D class average depicts 44 subunits in the top view. Each circle represents one subunit, with the starting one in red and the ending one in white.

Manually selected particles were used to train multiple Topaz models^17^. To minimize false positive selections, multiple rounds of 2D classification were carried out. If selected particles were separated by less than 16 nm (half the cage diameter) they were treated as duplicates and removed. An ab-initio reference map was generated with the remaining 3,730 particles, without symmetry constraints applied, and refined to ~ 16 Å (Fig. S2).

Two-dimensional classification pointed to rotational symmetry (Fig. 2C), and in one class there appeared to be 44 subunits (Fig. 2D). A rotational symmetry search from C1 to C51 yielded the highest resolution with C44 symmetry (Figure S2). 3D classification without alignment followed by further refinement and local motion correction improved the resolution to 3.5 Å. Still, some regions of the map, particularly at the ends of the truncated cone, were not well resolved (Fig. S3A).

Two proteins, Flotillin1 and Flotillin2, which are 69% identical, were identified in Western blot analysis of vesicles isolated from HEK293 GnTl^−^ cells (supplementary table 1) along with several other members of the Stomatin, Prohibitin, Flotillin, HflK/C (SPFH) family^3^. The structure of one member of this family, HflK/C from bacteria, forms a cage that attaches to membranes^14^. We hypothesized that the Flotillins might form the larger cages in our samples. Past studies showed that Flotillin1 and Flotillin2 co-localize in puncta^6^, that knockdown of either one decreases expression of the other^18^, and that the two can be cross-linked^4^. We therefore considered the possibility that both Flotillins form a pseudo-symmetric structure. We applied C22 symmetry with a mask over the poorly defined narrow end of the truncated cone. This procedure distinguished between two layers of helical density (Figure S3C). One layer contains 44 ‘cap’ helices and the other 22 ‘CTD’ helices, explicable if one Flotillin encodes a CTD helix and the other does not. We conclude that the cage consists predominantly of 22 each of Flotillin1 and Flotillin2 in an alternating pattern.

C22 symmetry applied to the entire structure improved density in many regions. For example, residues R291 of Flotillin 1 and A294 of Flotillin2 are poorly resolved with C44 symmetry but improved with C22 symmetry (Fig. S3D). Similar improvements in side-chain density were observed for corresponding residues in Flotillin1/Flotillin2, including K360/A363, D355/A358, A339/K342, and F325/A328 (Fig. S3E). With a final density map for the overall Flotillin complex at 3.5 Å resolution, atomic models could be built for much of the molecule. Regions with poor or weak density were removed. The final model consists of Flotillin-1:1-422 and Flotillin-2: 1-402. (Fig. S4A), except for the SPFH1 domains (Fig. S4B). The density extends from the 44-helical layer to the 22-helical layer in every other subunit, permitting us to identify the origin of the 22-helical layer of Flotillin1 (Fig. S4C). The segment following the cap-helix domain in both Flotillin1 and Flotillin2 shows some differences. For example, the length of the linker connecting the cap helix to the CTD helix aligns with the sequence of Flotillin1. The model is consistent with constraints derived through lysine-lysine cross-link data^4^. Thus, it is likely that 22 pairs of Flotillin 1 and Flotillin 2 form the cage, however, we cannot exclude the possibility that other stoichiometries can occur.

The Flotillin protein complex resembles a basket, with its wide end always connected to a membrane and its narrow end sometimes connected to a separate membrane (Fig. 3A). The wide end has a diameter of 32 nm, the narrow end 19 nm, and height 22 nm (Fig. 3B). The wide end contains SPFH1 and SPFH2 domains at the N-terminus (Fig. 3C). The wall is a single layered α-helical barrel, and the narrow end consists of the two α-helical layers described above, one with 22 and the other with 44 α-helices (Fig 3D). A β-strand from each Flotillin makes a 44-stranded β-barrel inside the helical layer. C-terminal amino acids form a poorly resolved structure that further constricts the opening of the narrow end to about 4 nm diameter.

**Fig 3.**
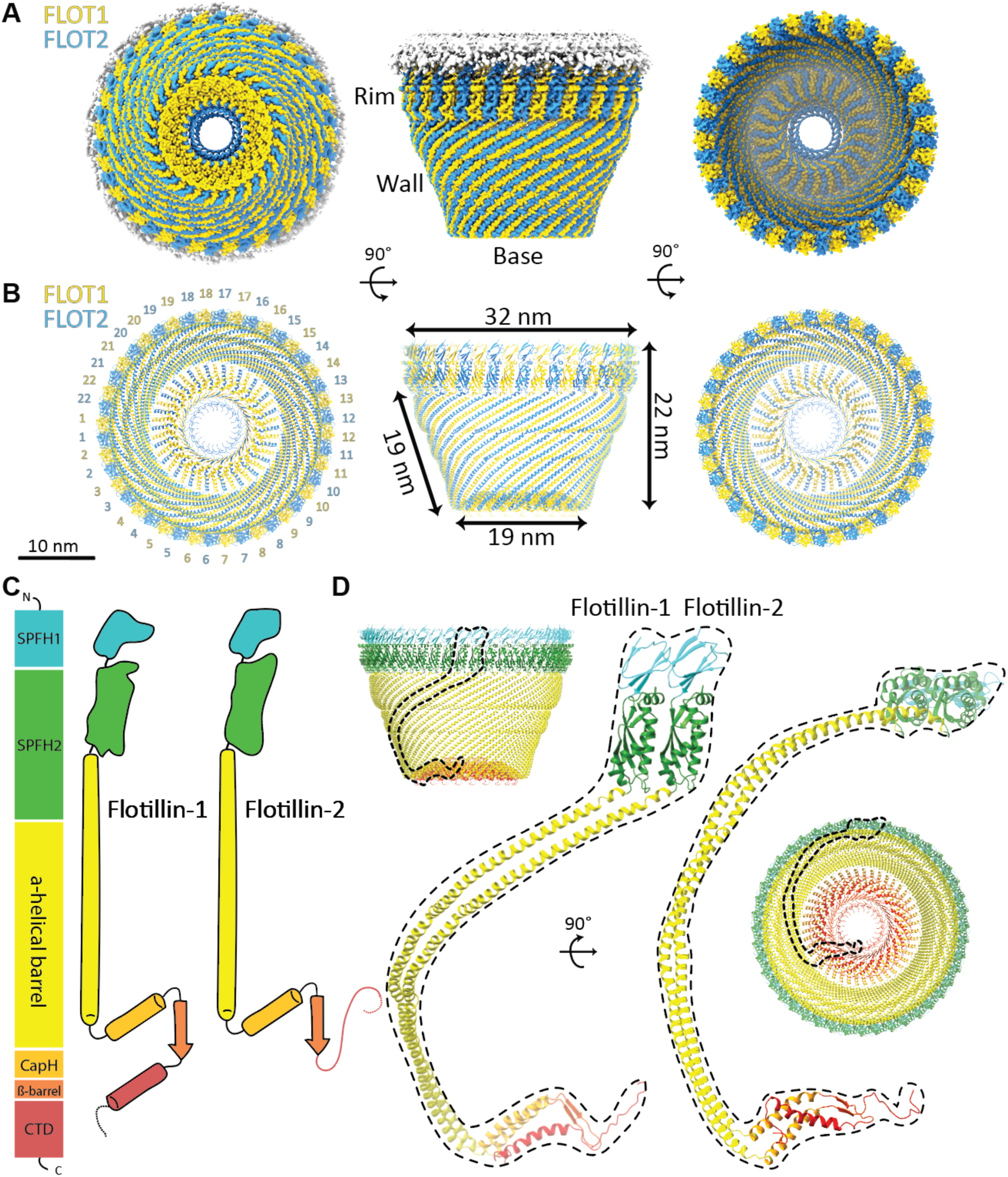
The Overall Structure of the Flotillin Complex and Domain Organization. **(A)** Views from the bottom (cytoplasmic view), side, and top of the cryo-EM map of the Flotillin complex. Flotillin1 and Flotillin2 are denoted in yellow and blue, respectively, associated with white-shaded cryo-EM density corresponding to the membrane. Regions corresponding to the Rim, Wall, and Base are labeled adjacent to their respective locations in the cryo-EM map. **(B)** Views from the bottom (cytoplasmic view), side, and top of the atomic model representing the Flotillin complex. Labels accompany each part, providing dimensions for the Flotillin complex. **(C)** Schematic depiction illustrating the domain organization and secondary structure topology of Flotillin1 and Flotillin2. Individual domains are colored: SPFH1 in cyan, SPFH2 in green, Flotillin α-helical barrel in yellow, cap helix (CapH) in orange, β-barrel in orange red and C-terminal domain (CTD) in red. Flotillin1 and Flotillin2 differ in their CTD domains. **(D)** The atomic model of the dimer of Flotillin1 and Flotillin2. Secondary structural motifs are color-coded as described in (C). Dashed lines delineate the dimer’s positions relative to the entire Flotillin complex from both side and bottom views.

### Structural Basis of Flotillin Oligomerization

The C-terminus of Flotillin was suggested to be involved in oligomerization^19^. The base structure is made by the C-terminus of Flotillin arranged in two layers of helices that run in opposite directions and are linked by a β-strand (Fig. 4A and B). From the structure, we observed strong interactions between Flotillin1 and Flotillin2 in the base structure.The first layer has 44 helices from both Flotillin1 and 2 (22 each). The interaction between Flotillin1 and Flotillin2 is enhanced by ionized hydrogen bonds (Fig. 4C and D). The center of the base is a parallel β-barrel containing 44 β-strands from both Flotillin1 and 2. After the β-barrel, Flotillin1 and Flotillin2 diverge. Flotillin1 extends outward, while Flotillin2 goes further in towards the center. Flotillin1 adds a new layer of 22 helices on top of the first layer, placing hydrophobic residues between the two helical layers. (Fig 4E).

**Fig 4.**
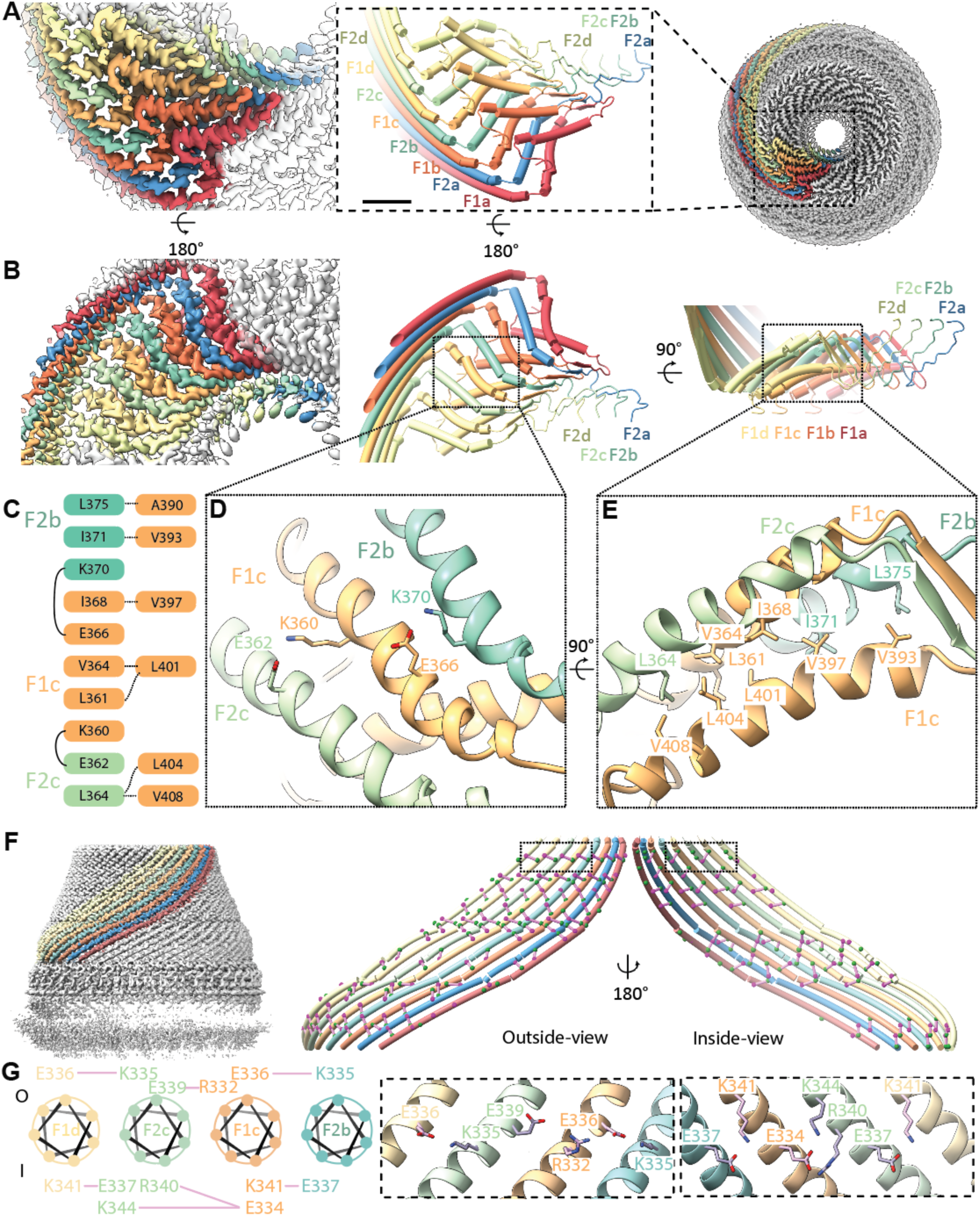
Structure and Organization of the Base and Wall Regions in the Flotillin Complex. **(A)** Bottom view (right) of the Flotillin complex displaying eight subunits, each uniquely colored. In the middle, a zoomed-in view of the boxed region highlights the cartoon representation of the Base, consisting of eight subunits (F1a, F2a, F1b, F2b, F1c, F2c, F1d, F2d), with their corresponding cryo-EM density shown on the left. **(B)** Top view of the Flotillin complex’s cryo-EM density (left) and cartoon representation (middle). On the right, the side view of the same Base region is presented. **(C)** Amino acids involved in subunit interactions, with colors indicating amino acids from the same subunits. Dashed lines represent hydrophobic interactions between the first (left) and second (right) helix-layers, while solid lines indicate inter-subunit ionized hydrogen bonds. **(D)** Zoomed-in top view detailing interactions in the Base highlighting the ionized hydrogen bonds. **(E)** Zoomed-in side view detailing interactions in the Base highlighting close contacts that would permit van der Waals interaction. **(F)** Side view of the Flotillin complex with eight Wall helices separately colored on the left. On the right, both outside-view and inside-view of the eight Wall helices highlight charged residues involved in forming ionized hydrogen bonds (colored in green and magenta), with interactions represented by solid purple lines connecting the α-carbons of interacting amino acids. **(G)** Wheel plot illustrating amino acids from four neighboring subunits involved in inter-subunit ionic hydrogen bonds. The top part shows residues outside the Flotillin complex (labeled as O), and the bottom part shows inside residues (labeled as I). Dashed box regions correspond to the zoomed-in view from (F), displaying detailed interactions between charged amino acids involved in ionic hydrogen bonding.

The wall of the Flotillin cage is made of a right-handed α-helical barrel with 44 helices arranged side by side (Fig. 4F). This formation spans 186 amino acids, constituting the core of the Flotillin structure. These helices are characterized by repeating patterns of amino acids (A/G, E, A/G, E), which are believed to form coiled-coil bundles that facilitate the clustering of Flotillin molecules^1^. Flotillin does not contain heptad-repeat sequences found in classical coiled-coil helices. Instead, Flotillin is characterized by the presence of charged amino acids both inside and outside its α-helical barrel. These charged residues form specific clusters of ionized hydrogen bonds, likely serving to glue adjacent subunits together (Fig 4F and G and Fig. S5).

The structure of Flotillin’s right-handed helical barrel differs from those found in other proteins like SNARE^20^ and motor proteins such as myosin, kinesin, and dynein^21^. These proteins typically have helices that intertwine in pairs or groups, unlike Flotillin’s parallel arrangement. This difference notwithstanding, the buried area per residue in Flotillin’s wall structure is like that in these other helical structures (detailed in the methods section).

### Interactions Between Flotillin and Membranes

In our micrographs the wide end of the Flotillin cage is mostly bound to a membrane surface. Adherence to the membrane is mediated by the SPFH1 domains, which penetrate one leaflet of the lipid bilayer (Fig. 5A). We show in Figure 5B that the SPFH1 domains expose to the membrane a surface containing many hydrophobic amino acids (Fig. 5B). We note the prevalence of Trp residues, which are well known to be most stable near the membrane’s headgroup layer (Fig. 5B, solid line). We also note the many cysteine residues on the hydrophobic surface. Some of these are palmitoylated^19^, and Flotillin-2 is N-myristoylated on Gly2. This myristoylation is required for membrane association^19^. The bacterial HflK/HflC complex attaches to membranes through transmembrane helices^14^ (Ma 2022). The Flotillin cage, by contrast, uses a hydrophobic surface that contains covalent lipid molecules.

**Fig 5.**
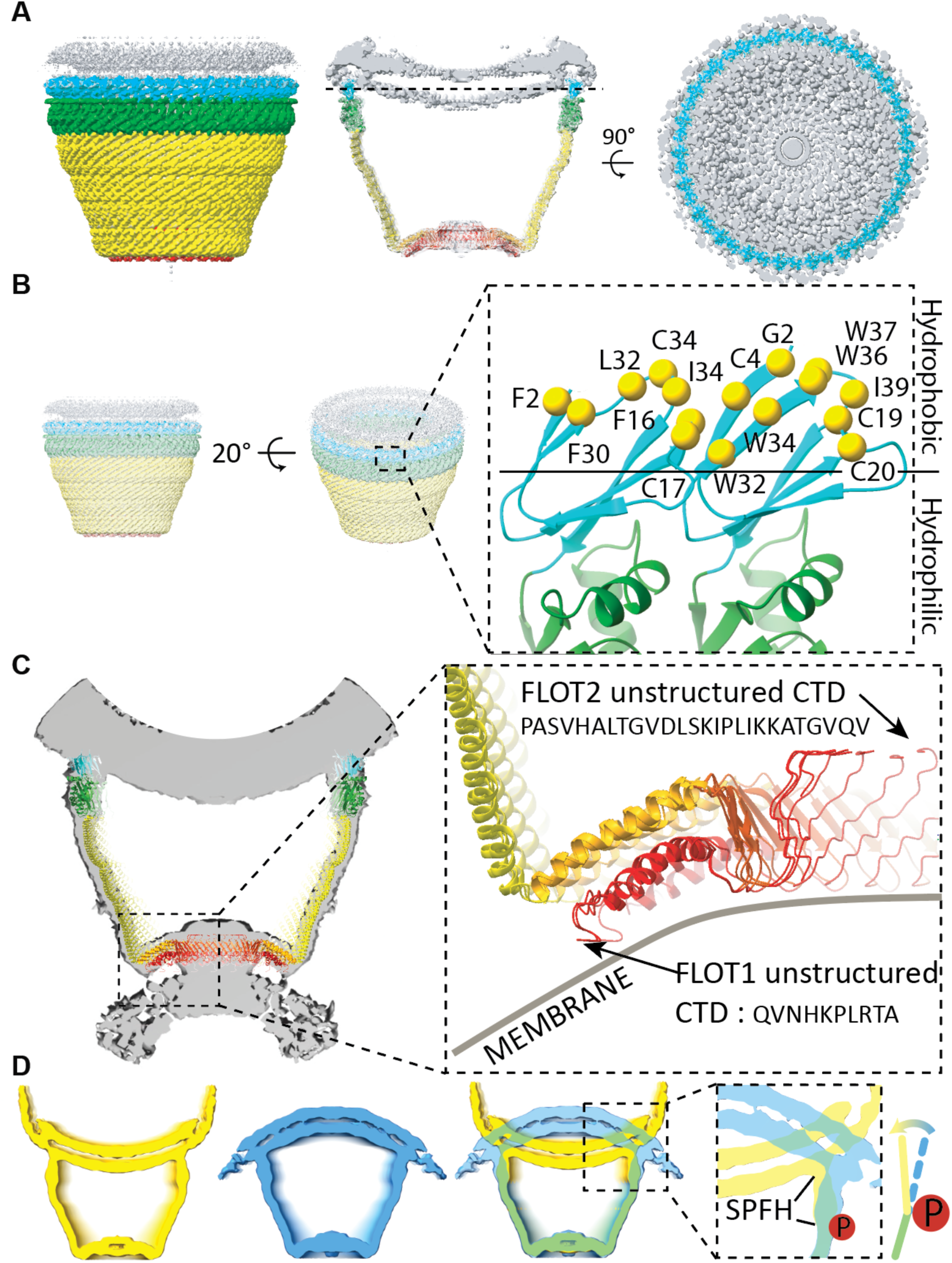
Membrane Association of the Flotillin Complex. **(A)** Side view (left), intersection view (middle), and plasma membrane intersection view at the plane indicated by the dashed line from the middle panel. Individual domains are distinctly colored: SPFH1 in cyan, SPFH2 in green, CC1 in yellow, CC2 in orange, and CTD in red. The membrane is depicted in grey. **(B)** Zoomed-in view of the SPFH1 domain from the dimer of Flotillin1 and Flotillin2 cartoon representation (right), as indicated by the dashed boxed region from the 20-degree tilted side view on the left. Hydrophobic residues and potential lipidated cysteines, halfway inserted in the inner leaflet of the membrane (above the solid line), are highlighted in yellow and labeled accordingly. **(C)** Reconstruction of the Flotillin complex associated with two membranes. The dashed box region highlights potential parts involved in membrane association in the Base region. The sequence of the unstructured region of Flotillin 1 and Flotillin 2 CTDs is shown, with an arrow indicating the beginning of the unstructured region at CTDs. Membrane positions are depicted with grey lines and labeled. **(D)** Intersection view of cryo-EM maps of the Flotillin complex outside (in yellow) and inside (blue) vesicles. An overlapping view of both cryo-EM maps with a focus on the conformational change in the SPFH domain (dashed box). The bending motion of SPFH domain in the Flotillin complex is indicated using colored lines and a directional arrow. The potential site of phosphorylation is marked by a filled red circle containing the letter ‘P’.

When Flotillin cages were located outside a vesicle, they often connected to a second membrane at their narrow end (Fig. 2B). The unstructured C-terminus regions of Flotillin1 and Flotillin2 are likely involved in this membrane interaction, although the low resolution of this region precludes a detailed description (Fig. 5C).

Flotillin cages in outside-out and inside-out orientations showed conformational differences at the membrane interface (Fig. 5D), where inward bending of the SPFH domains may induce membrane invagination and initiate a clathrin-independent endocytic pathway^6^. This bending may be initiated by the phosphorylation of tyrosine residues Tyr160 in Flotillin1 and Tyr163 in Flotillin2 by kinase Fyn. Located at the hinge region, phosphorylation of Tyr160 and Tyr163 is essential for altering membrane curvature and initiating endocytosis, as demonstrated by the reduced endocytosis observed when these residues are mutated to phenylalanine^7^. Further research is needed to explore how these phosphorylation-induced changes impact Flotillin’s functionality in endocytosis and membrane dynamics.

## Discussion

### Structural insights for SPFH protein family

The SPFH protein family members likely share a similar architecture. Using negative stain cryo-EM Prohibitin, Slr1128/SYNY3, and Erlin-1/2 were all shown to form ring-shaped structures^9,22–24^. More recently, the aforementioned HflKC and FtsH complex was determined at high resolution^14,25^.

Using Alphafold2^26^, we predicted SPFH proteins’ hetero/homo-tetramer structures based on their sequences from SPFH2 to the C-terminal cap helix which they share in common (Fig. 6A). Different SPFH family proteins have various lengths of wall helices ranging from 189 to 44 amino acids. Following the long helix, they share a similar cap helix forming the first helical layer, resembling the narrow end of the Flotillin cage. The differing lengths of the wall helix and cap helix likely influence the overall size and the number of subunits in their protein complexes (Fig. 6B). Additionally, we observed that two other truncated-cone structures in the native membrane vesicles of HEK293 GnTl^−^ cells, presumably stomatin and stomatin-like protein 2, exhibit distinct sizes and shapes compared to the Flotillin complex (Fig. 6C). This suggests that SPFH family proteins can form a variety of cage structures.

**Fig 6.**
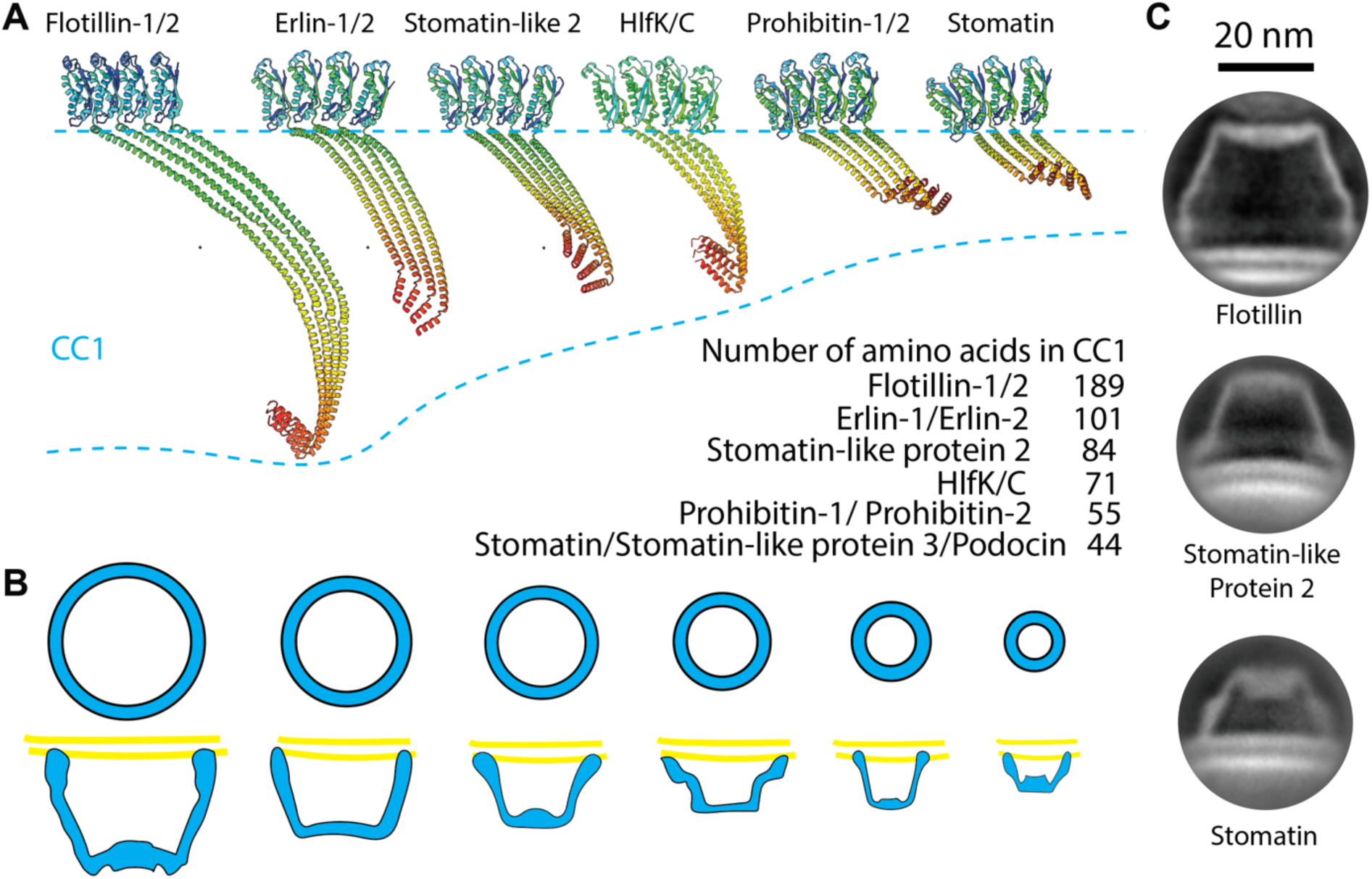
Common Architecture of the SPFH Protein Family and Their Functional Implications. **(A)** Predicted tetrameric protein complexes of Flotillin1/2, Erlin1/2, Stomatin-like protein 2 (SLP2), HlfK/C, Prohibitin1/2, and Stomatin by Alphafold2. The complexes contain SPFH2, CC1, and CC2 domains, with monomers color-coded from green to red representing N-terminus to C-terminus. The table lists the number of amino acids in the CC1 domain of each protein. **(B)** General model illustrating the formation of a basket-like structure associated with the membrane by SPFH proteins. Varying lengths of SPFH proteins result in differences in the area they occupy on the membrane and the size of their complexes. **(C)** Representative cryo-EM 2D classification averages of the Flotillin complex, presumably Stomatin-like protein 2, and Stomatin determined from native membrane vesicle preparations. The scale bar is 20 nm.

### Roles of Flotillin

Flotillin is mainly found on the surface of cells, at the plasma membrane, but it also connects with membranes inside cell compartments such as exosomes^27^, endosomes^28^, lysosomes^10^, and the nucleus^29^. The presence of Flotillin in these various locations suggests its role in regulating multiple cellular processes^27,30,31^.

Flotillin1 and Flotillin2 are conserved proteins found across various species, with some possessing additional Flotillin-like proteins^32,33^. These variants contain flotillin domains necessary to form cage complexes and also contain functional domains tethered at their N-terminus, C-terminus, or both. These tethered functional domains exhibit a wide range of functions, including DNA/RNA-interactions^34,35^, nucleotide binding^36^, gene regulation^37^, metabolic processes^38,39^, immune responses^40^ and protein transport^41^ (Figure S6). The spatial organization of functional domains relative to the Flotillin cage—whether inside or outside—might depend on their functional requirements. For example, domains like glycoside hydrolase that need protection may be located inside the cage for regulated activity, while domains requiring interaction freedom, such as kinesin motors, are likely positioned outside^42^. This structural arrangement of the Flotillin complex allows for diverse functions and spatial control over these activities.

From an evolutionary perspective, tethering functional domains to Flotillin enhances cellular functionality and adaptation, a strategy also common in other SPFH family proteins. This suggests a widespread evolutionary approach, even when proteins are not directly tethered^14^. In Methicillin-resistant Staphylococcus aureus (MRSA), Flotillin facilitates the clustering of PBP2a, a protein crucial for penicillin resistance. Alterations in Flotillin disrupt this clustering, reducing MRSA’s resistance^43^.

The finding in this study that Flotillin cages locally alter membrane curvature is intriguing. By exerting bending forces that act locally, Flotillin might influence mechanical properties of membranes and thus play a role in mechanosensory channel function. This idea is supported by studies in round worms and mice, where Mec-2 and Stomatin-like protein 3 (similar to Flotillin) are essential for touch sensitivity^44–46^. It has been hypothesized that these proteins may confine mechanosensory channels within their cage structures to regulate them^47^. Further research will be needed to confirm whether mechanosensitive channels collaborate with SPFH proteins or if other mechanisms are at play^48–50^.

In summary, we demonstrated the use of various less disruptive preparations to preserve membrane and membrane associated proteins from different cells. From these preparations, we determined a fascinating protein complex structure, the Flotillin cage, using cryo-EM and single-particle analysis. We expect that more interesting discoveries will be made, and more exciting biological questions raised through these more native structural approaches.

## Materials and Methods

### Cell Culture

HEK293S GnTl^−^ cells (ATCC CRL-3022) were cultured in Freestyle 293 medium supplemented with 2% fetal bovine serum at 37 °C. A431 cells (ATCC CRL-1555) were cultured in Dulbecco’s Modified Eagle’s Medium, Catalog No. 30-2002 supplemented with 10% fetal bovine serum. Human erythrocytes were purchased from Innovative Research (IWB3ALS40ML).

### Plasma Membrane Vesicle Preparation

Four liters of live HEK293 GnTl^−^ cells (3 × 10^6^/mL) were harvested by centrifugation at 500 g for 10 min. The cell pellet was resuspended in 500 mL GPMV buffer containing 10 mM K-HEPES, pH 7.4, 140 mM NaCl, 10 mM KCl, and 2 mM CaCl2. The cells were centrifuged again at 500 g for 10 min and then resuspended in 500 mL GPMV buffer supplemented with NEM (Thermo Scientific™ Pierce™, 23030) at 7.5 mM. The cell resuspension was transferred to four 500 mL baffled flasks to contain 125 mL in each flask. The flasks were then incubated in a 37 °C incubator shaking at 130 rpm for 2 h. The fast shaking helps to dislodge the native membrane vesicles from the cells, resulting in smaller unilamellar vesicles. At the end of the incubation, the flasks were shaken by hand for ~30 s. The suspension was then spun down at 500 g at 4 °C for 10 min to remove cells and large vesicles. Next, 10% glycerol was added to the supernatant to prevent aggregation, and it was sonicated using a Branson probe sonicator (1/2” tip with Branson102-C converter) at 40% power in three pulses of 30 seconds each, with approximately one minute of cooling on ice between each pulse. The vesicles were then pelleted by ultracentrifugation at 100,000 g at 4 °C for 1 h in a Ti70 rotor.

The membrane vesicle pellet from every ~25 mL sample was triturated in ~0.2 mL GPMV buffer supplemented with 10% glycerol by gently squirting the buffer toward the pellet (with a 100 μL tip). The resuspended vesicles in the ultracentrifugation tubes were then sonicated in a water bath sonicator (Branson M1800) with ~10 s pulses at room temperature until the solution was opalescent. The vesicles were subsequently spun down at 3,500 g at 4 °C for 10 minutes to remove any remaining aggregates.

The supernatant was mixed with two volumes of 90% sucrose GPMV buffer and loaded at the bottom of a step gradient that contained 2 ml of 45%(w/v) and 35% sucrose, 4ml of 25% and 12% sucrose GPMV solutions in 14-ml ultra-clear tube. Next the tubes were centrifuged at 150,000×g for 18 h at 4°C in a SW40Ti rotor. After centrifugation, 25%-45% fraction was collected and then concentrated to OD280 ~2 using an Amicon 2 mL concentrator (molecular weight cutoff of 100 kDa).

For Western blot analysis, vesicles were loaded onto sodium dodecyl sulfate– polyacrylamide gel electrophoresis (SDS-PAGE) using Bio-Rad Mini-PROTEAN® TGX™ Precast Gels (4 to 15%) and subjected to electrophoresis. Following electrophoresis, the proteins were transferred to a nitrocellulose membrane. The blots were then blocked for 30 minutes at room temperature in TBS-T (25mM Tris-HCl pH 7.4, 0.9% NaCl, and 0.02% Tween-20) containing 4% dry milk. Subsequently, the membranes were probed with anti-Flotillin-1 antibody (Protein tech, 15571-1-AP), anti-Flotillin-2 antibodies (Protein tech, 28208-1-AP), Stomatin (Santa Cruz, sc-376869), Stomatin-like protein 2 (Abcam, ab191883) and detected using fluorescence.

### Exosomes Preparation

4 liters of conditioned medium was harvested from live HEK293 cultured cells (3 × 10^6^/mL). Cells were removed by centrifugation at 3000 ×g for 20 min followed by 10,000 ×g for 30 min. The supernatant was centrifuged at ~100,000 ×g for 1 hr in a Ti70 rotor.

The membrane vesicle pellet from every ~25 mL sample was triturated in ~0.2 mL GPMV buffer supplemented with 10% glycerol by gently squirting the buffer toward the pellet (with a 100 μL tip). The resuspended vesicles in the ultracentrifugation tubes were then sonicated in a water bath sonicator (Branson M1800) with ~10 s pulses at room temperature until the solution was opalescent. The vesicles were then centrifuged at 3,500 g at 4 °C for 10 min to remove the remaining aggregates.

The supernatant was mixed with two volumes 90% sucrose GPMV buffer and loaded at the bottom of a step gradient that contained 2 ml of 45%(w/v) and 35% sucrose, 4ml of 25% and 12% sucrose GPMV solutions in 14-ml ultra-clear tube. Then, the tubes were centrifuged at 150,000×g for 18 h at 4°C in a SW40Ti rotor. After centrifugation, 25%-45% fraction was collected and then concentrated to an absorbance at 280 nm of ~2.0 using an Amicon 2 mL concentrator (molecular weight cutoff of 100 kDa).

### Preparation of Total Membrane Vesicles

All procedures were conducted at 4 °C within a cold room. Pellet from 4 L of cells was resuspended in 160 mL of lysis buffer (20 mM K-HEPES pH 7.4, 300 mM KCl, 0.5 mM MgCl2, and 5 mM dithiothreitol (DTT)) supplemented with 10 µg/mL leupeptin, 10 µg/mL pepstatin A, 1 mM benzamidine, 2 μg/mL aprotinin, 0.3 mg/mL AEBSF, ~100 µg/mL DNase, and ~100 µg/mL RNase A. Then the supernatant was centrifuged at ~100,000 × g for 1 hr in a Ti70 rotor. The membrane vesicle pellet from every ~25 mL sample was triturated in ~0.2 mL GPMV buffer supplemented with 10% glycerol by gently squirting the buffer toward the pellet (with a 100 μL tip). The resuspended vesicles in the ultracentrifugation tubes were then sonicated in a water bath sonicator (Branson M1800) with ~10 s pulses at room temperature until the solution was opalescent. The vesicles were then centrifuged at 3,500 g at 4 °C for 10 min to remove the remaining aggregates. The supernatant was mixed with two volumes 90% sucrose GPMV buffer and loaded at the bottom of a step gradient that contained 2 ml of 45%(w/v) and 35% sucrose, 4ml of 25% and 12% sucrose GPMV solutions in 14-ml ultra-clear tube. Then the tubes were centrifuged at 150,000×g for 18 h at 4°C in a SW40Ti rotor. After centrifugation, 25%-45% fraction was collected and then concentrated to OD280 ~2 using an Amicon 2 mL concentrator (molecular weight cutoff of 100 kDa).

### Erythrocytes Vesicle Preparation

Washed human erythrocytes (Innovative Research, IWB3ALS40ML) were hemolyzed in 5 volumes of 1 mM EDTA and 10 mM Tris-HCl, pH 7.4. Then they were centrifuged at 18,000 g for 10 min. The resulting ‘ghost’ membranes were then washed five times in the hemolysis buffer with 1 mM EDTA and 10 mM Tris-HCl, pH 7.4. Ghost membranes were pelleted (18,000 g, 10 min) and then sonicated using a probe sonicator (Branson, 1/2” tip with Branson102-C converter) at 60% power for three 30 s pulses with ~1 min chilling on ice in between pulses. The vesicles were then centrifuged at 18,000 g at 4 °C for 10 min twice to remove the remaining aggregates. The vesicles were then pelleted by ultracentrifugation at 100,000 g at 4 °C for 2 h in a Ti70 rotor.

The membrane vesicle pellet from every ~25 mL sample was triturated in ~0.2 mL of GPMV buffer supplemented with 10% glycerol by gently squirting the buffer toward the pellet (with a 100 μL tip). The resuspended vesicles in the ultracentrifugation tubes were then sonicated in a water bath sonicator (Branson M1800) with ~10 s pulses at room temperature until the solution was opalescent. The vesicles were then centrifuged at 3,500 g at 4 °C for 10 min to remove the remaining aggregates.

The supernatant was mixed with two volumes 90% sucrose GPMV buffer and loaded at the bottom of a step gradient that contained 2 ml of 45%(w/v) and 35% sucrose, 4ml of 25% and 12% sucrose GPMV solutions in 14-ml ultra-clear tube. The tubes were then centrifuged at 150,000 × g for 18 h at 4°C in a SW40Ti rotor. After centrifugation, the 25%-45% fraction was collected and concentrated to an absorbance at 280 nm ~2.0 using an Amicon 2 mL concentrator (molecular weight cutoff of 100 kDa).

### Cryo-EM Grid Preparation for Membrane Vesicles

Quantifoil R1.2/1.3 400 mesh holey carbon gold grids were glow-discharged for 22 s. Then 3 μL of concentrated membrane vesicles were applied to freshly glow-discharged grids and left for 10 min at 22 °C with a humidity of 100%. The grids were then manually blotted at the edges using a piece of filter paper. Another 3 μL sample was applied, and grids were blotted using a Vitrobot Mark IV with a blot force of 1 for 5 seconds following a 30-second incubation. The grids were quickly frozen in liquid ethane and kept in liquid nitrogen until data collection.

### Cryo-EM Grid Preparation for Unroofed Erythrocytes

Washed human erythrocytes (Innovative Research, IWB3ALS40ML) were diluted at a ratio of 1:100 and deposited onto carbon-coated 200-mesh copper grids. These grids had previously undergone glow discharge and were pre-coated with a poly-D-lysine solution. Following a 30-minute incubation, the human erythrocytes were rinsed in a low salt buffer (a 1:3 dilution of Dulbecco’s phosphate-buffered saline (DPBS)). The unroofing process involved the application of a stream of DPBS through a 20-gauge needle. Subsequently, the unroofed erythrocytes were washed three times at room temperature with DPBS.

3 μL DPBS was then applied to the grids containing the unroofed erythrocytes. The grids were manually blotted from the edge using a piece of filter paper and were plunge-frozen using a Vitrobot Mark IV. The blotting was performed with a blot force of 0 and a blot time of 3 seconds.

### Cryo-electron Tomography Data Collection and Data Processing

Tilt series were collected using a Titan Krios (Thermo Fisher) using a Gatan Bioquantum energy filter and a K3 direct detector at 300 keV. Data collection was symmetric with a tilt range from −42° to 42° in 3° steps, using SerialEM. Each tilt image was recorded in 400-ms frames at an approximate defocus of 4 μm. The total dose per tilt series collected was 103 e /Å^2^, with dose rates of approximately 20 e /pixel per second at a pixel size of 2.6 Å. Full-frame alignment was performed using MotionCor2^51^.

Tilt series were aligned using Appion-Protomo^52^. Tilt series were initially aligned and then manual adjusted. The optimal iteration was reconstructed for visual inspection using the Tomo3D simultaneous iterative reconstruction technique^53^, with dose compensation applied according to a method previously reported^54^. The reconstructed tomograms were visualized using 3DMOD.

### Cryo-EM of Native Membrane Vesicles from HEK293 GnTl^−^ Cells

Cryo-EM data for native membrane vesicles from HEK293 GnTl^−^ cells were collected using a 300 keV Titan Krios transmission electron microscope with a cold-field emission gun and a 6 eV energy filter. A total of 19,024 micrographs were obtained by combining two datasets from sessions spanning six days. The microscope settings remained consistent throughout.

Considering the relatively low natural abundance of the Flotillin complex in plasma membrane vesicles, our estimation, based on preliminary screening results with a lower-end microscope, suggested one Flotillin complex image per 24 micrographs with a 4K x 4K camera at a pixel size of 1.5 Å. To reach a compromise between achievable resolution and the number of images of the Flotillin complex, we opted to collect data at a pixel size of 1.196 Å. The data collection pattern employed was a 3×3 grid with 2 shots per hole using SerialEM and a defocus range of −1.0 to −2.0 μm. The movies have a total dose of 42 e^−^/Å^2^.

Raw movies were motion-corrected and CTF parameters estimated using patch-based methods in cryoSPARC (V4.2.0)^55^. The initial 1,000 particles were manually picked to train multiple Topaz models for picking particles from all micrographs. Duplicate particles were excluded if they were less than 16 nm apart. Several rounds of 2D classification were then conducted to isolate particles with Flotillin density. The Flotillin model was initially generated using ab initio reconstruction with C1 symmetry. 3,730 particles were selected for further refinement. Particles are refined based on C1-C51 symmetry in different runs. Refinement results expected for C11, C22, C43 and C44 symmetry showed little helical feature. We conducted 3D classification without image alignment in Relion to eliminate particles with low-resolution contributions to the reconstruction^56^. One high-resolution class containing most particles (1,436) with C44 symmetry was selected for local motion correction in cryoSPARC. Then masked local refinement with C22 symmetry was applied to resolve the density near the narrow end of the Flotillin complex. This refinement yielded cryo-EM density at a resolution of 3.5 Å verified by the gold-standard FSC = 0.143 criterion. Local-resolution estimations were performed in cryoSPARC.

### Model Building, Refinement and Analysis

To build the Flotillin model in the high-resolution density map, the structure of the dimer of Flotillin-1 and Flotillin-2 predicted by AlphaFold was fitted into the map using molecular dynamic flexible fitting in ISOLDE^57^. We modelled four Flotillin dimers in the map to capture the entire interaction interface because a subunit interacts with multiple neighboring protomers in the region close to the narrow end. The four Flotillin dimer positions in the map were built manually in Coot^58^ and the other Flotillin subunits were adjusted in Coot by applying non-crystallographic symmetry, using the four Flotillin dimers as master chains. The structure underwent iterative refinement using Coot and Phenix real-space refine^59^ with non-crystallographic symmetry restraints while omitting geometry restraints. Regions with poor or weak density were removed. The final model consists of Flotillin-1:1-422 and Flotillin-2: 1-402. Refinement quality summary is in Supplementary Tables 2. Cryo-EM density maps and protein structures were visualized using UCSF ChimeraX^60^.

### Quantitative Analysis of Surface Area Contact Between Adjacent Protein Subunits

PDB structures (Vimentin: 3UF1, 104 aa; Myosin: 8G4L, 304 aa; Homer: 3CVE, 39 aa) were used for buried area per residue measurement. In ChimeraX, the command “measure buriedarea atom-spec1 withAtoms2 atom-spec2” was used to calculate buried area between two helices.

### Prediction of SPFH Family Protein Oligomeric Complex Structures by AlphaFold

The AlphaFold structures in this study were mainly generated from the AlphaFold2 implementation in the ColabFold notebooks^26^ running on Google Colaboratory^61^, using the default settings. For hetero/homo-tetrameric complex prediction, sequences were entered in tandem and separated by a semicolon. Due to computing memory constraints on Google Colaboratory, we used four protomers and removed the SPFH1 domain and CTD sequence. AlphaFold was run once with each of the five trained models; the five models generated were checked for consistency, and unless specified otherwise, the top-ranked model was taken in each case for density fitting. The AlphaFold predicted model was colored in rainbow from N-terminus to C-terminus and displayed in similar orientation as the Flotillin complex model.

### Data, Materials, and Software Availability

Cryo-EM density maps and atomic coordinates of the Flotillin have been deposited in the Electron Microscopy Data Bank under accession code EMD-44792 and in the Protein Data Bank under accession code 9BQ2.

## Acknowledgments

We thank Yi Chun Hsiung for aiding in cell culture, Tao Xiao, Chen Zhao, and George Vaisey for invaluable advice, and all members of the MacKinnon Lab and Chen Lab for insightful discussions. We thank Mark Ebrahim, Johanna Sotiris, and Honkit Ng at the Evelyn Gruss Lipper Cryo-EM Resource Center of Rockefeller University for their assistance with cryo-EM data collection. We also acknowledge Nicholas Spellmon, Rui Yan, Zhiheng Yu, and the team at the HHMI Janelia Cryo-EM Facility for their support. This work was supported by NIH Grant GM43949 (to R.M.), and R.M. is an Investigator in the Howard Hughes Medical Institute.

## Author contributions

Z.F. and R.M. conceptualized the research, with Z.F. carrying out the experiments. Data analysis was conducted by Z.F., and both authors contributed to writing the manuscript.

## Supporting Information

**Fig. S1.**
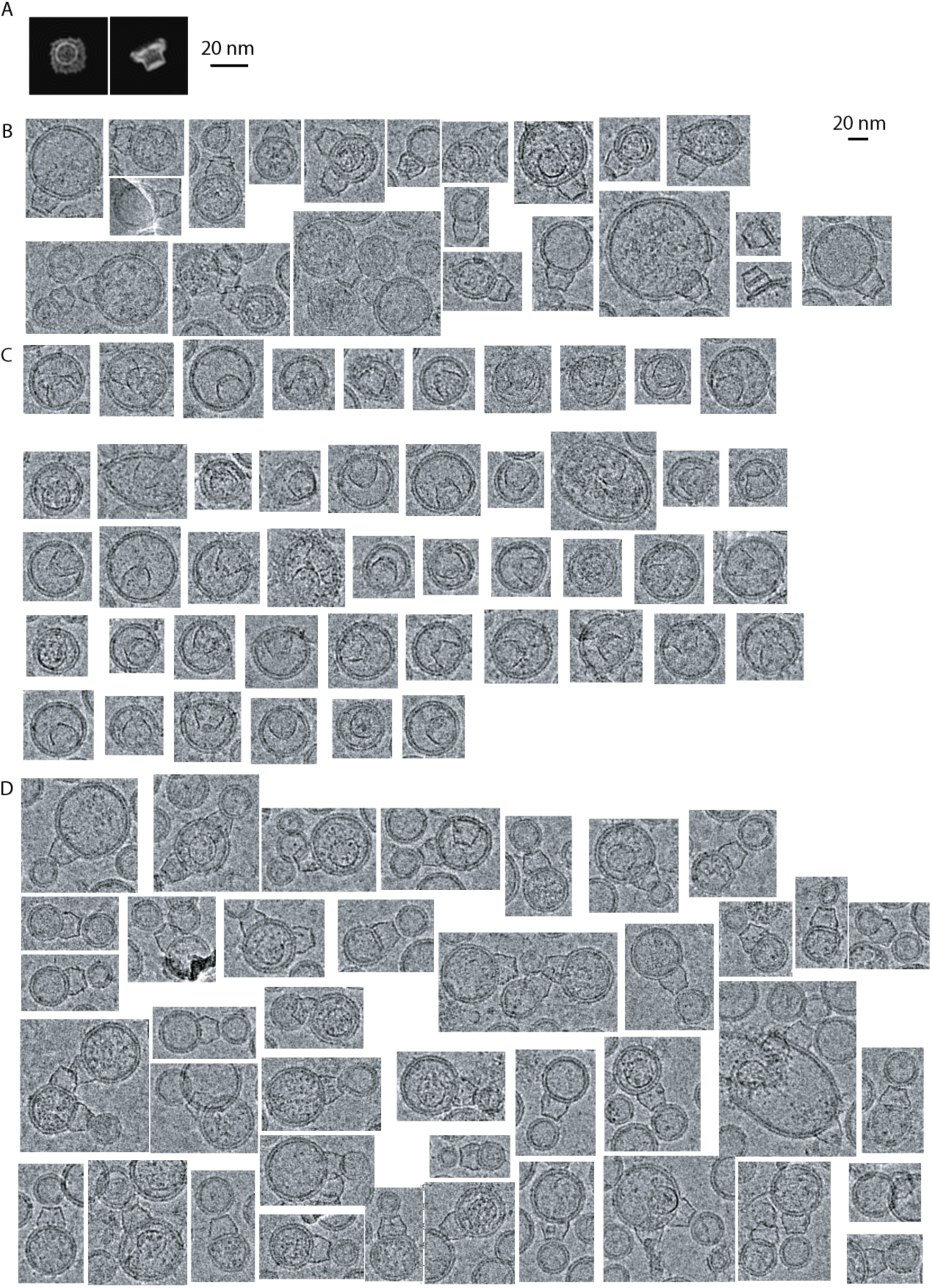
Dimension and Shape Comparison between the HlfK/C-FtsH Protein Complex and the Flotillin Complex. **(A)** Two-dimensional projections were generated from the cryo-electron microscopy map (EMD-32002) of the HlfK/C/FtsH protein complex. The projections were re-centered and lowpass filtered to 20Å. The scale bar represents 20 nm. **(B-D)** Exemplar cropped micrographs showcase the Flotillin complex observed ouside (B), inside (C), and bound to two (D) native membrane vesicles of HEK293 GnTl^−^ cells. The scale bar represents 20 nm.

**Fig S2.**
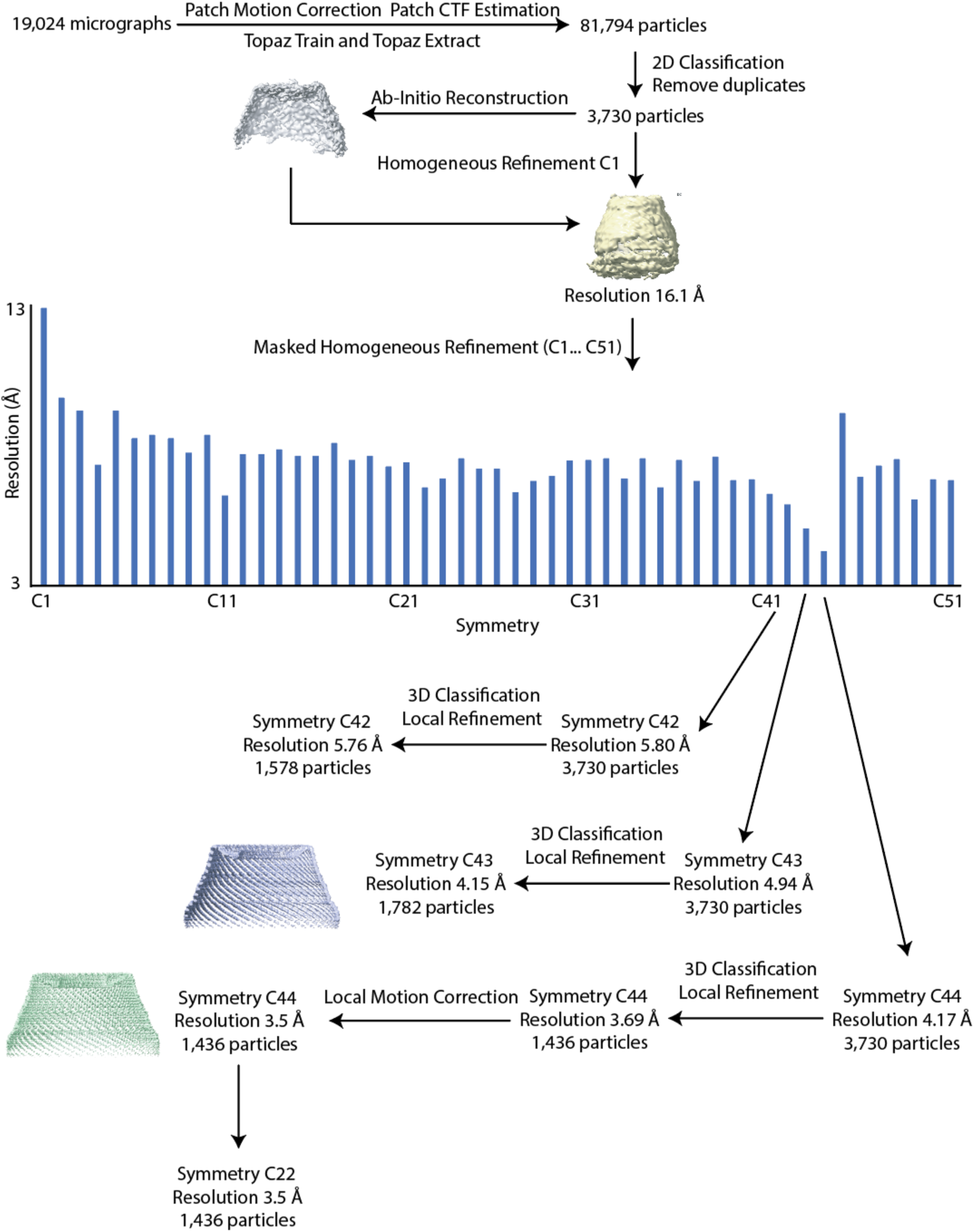
Cryo-EM data processing procedure.

**Fig. S3.**
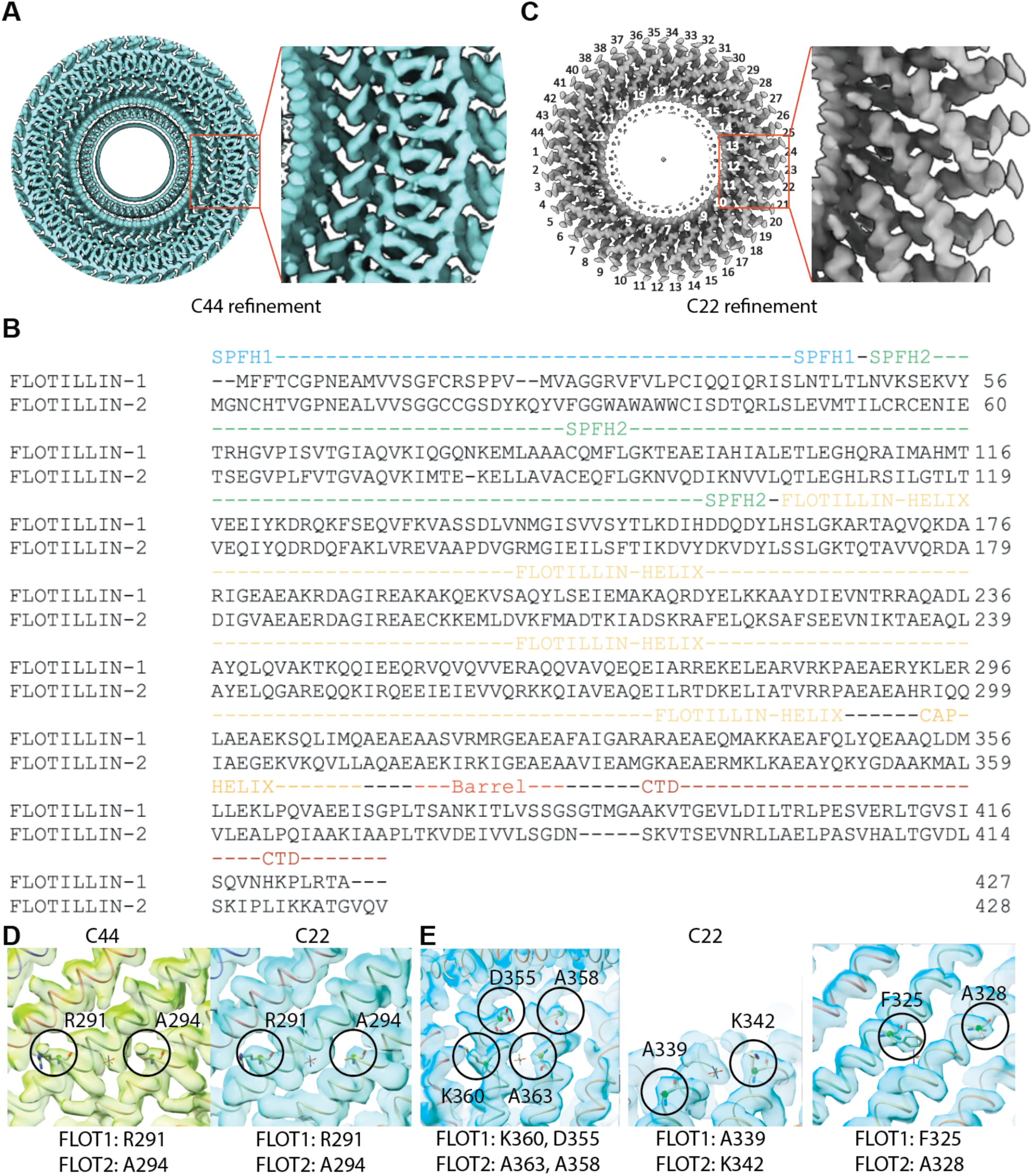
Symmetry of the Flotillin Complex. **(A)** The cryo-EM map of Flotillin’s narrow end region, reconstructed with C44 symmetry, exhibits a bottom view, revealing discontinuous density. **(B)** Sequence alignment of Flotillin-1 and Flotillin-2. **(C)** Bottom view of the Base of the Flotillin’s cryo-EM map reconstructed with C22 symmetry, showcasing well-resolved helical density. One layer comprises 44 helices (indicated with 1-44 in black), while the other layer contains 22 helices (indicated with 1-22 in white). **(D)** In the identical region of the cryo-EM map reconstructed with either C44 or C22 symmetry, the sidechain density becomes apparent, enabling the differentiation of Flotillin-1 and Flotillin-2. Circles emphasize the side chain exhibiting the most significant difference between Flotillin-1and Flotillin-2—specifically, R291 in Flotillin-1and A294 in Flotillin-2 at the corresponding position. The density, which was averaged out and fragmented under C44 symmetry, becomes distinctly resolved when employing C22 symmetry. **(E)** Representative regions of the cryo-EM map reconstructed with C22 symmetry, demonstrating distinct side chain density on the Flotillin-1and Flotillin-2 subunits.

**Fig. S4.**
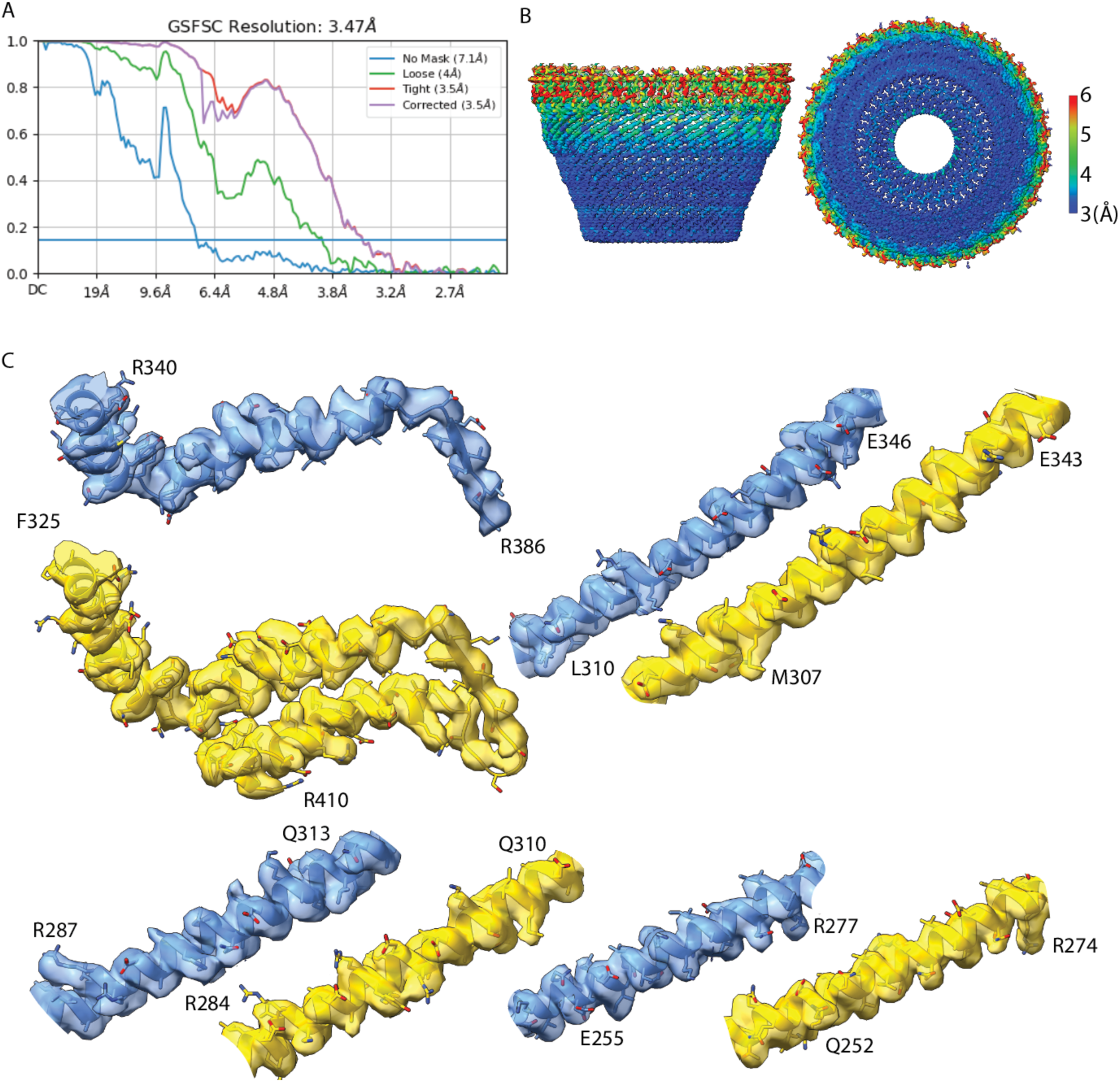
Resolution Estimation of the Final Density Map of the Flotillin Complex. **(A)** Fourier shell correlation (FSC) curves of the final map. At FSC gold-standard 0.143 cutoff, the estimated resolution is 3.5 Å. **(B)** Local resolution estimation of the final cryo-EM map. **(C)** Local densities of regions of Flotillin-1 (yellow) and Flotillin-2 (blue).

**Fig. S5.**
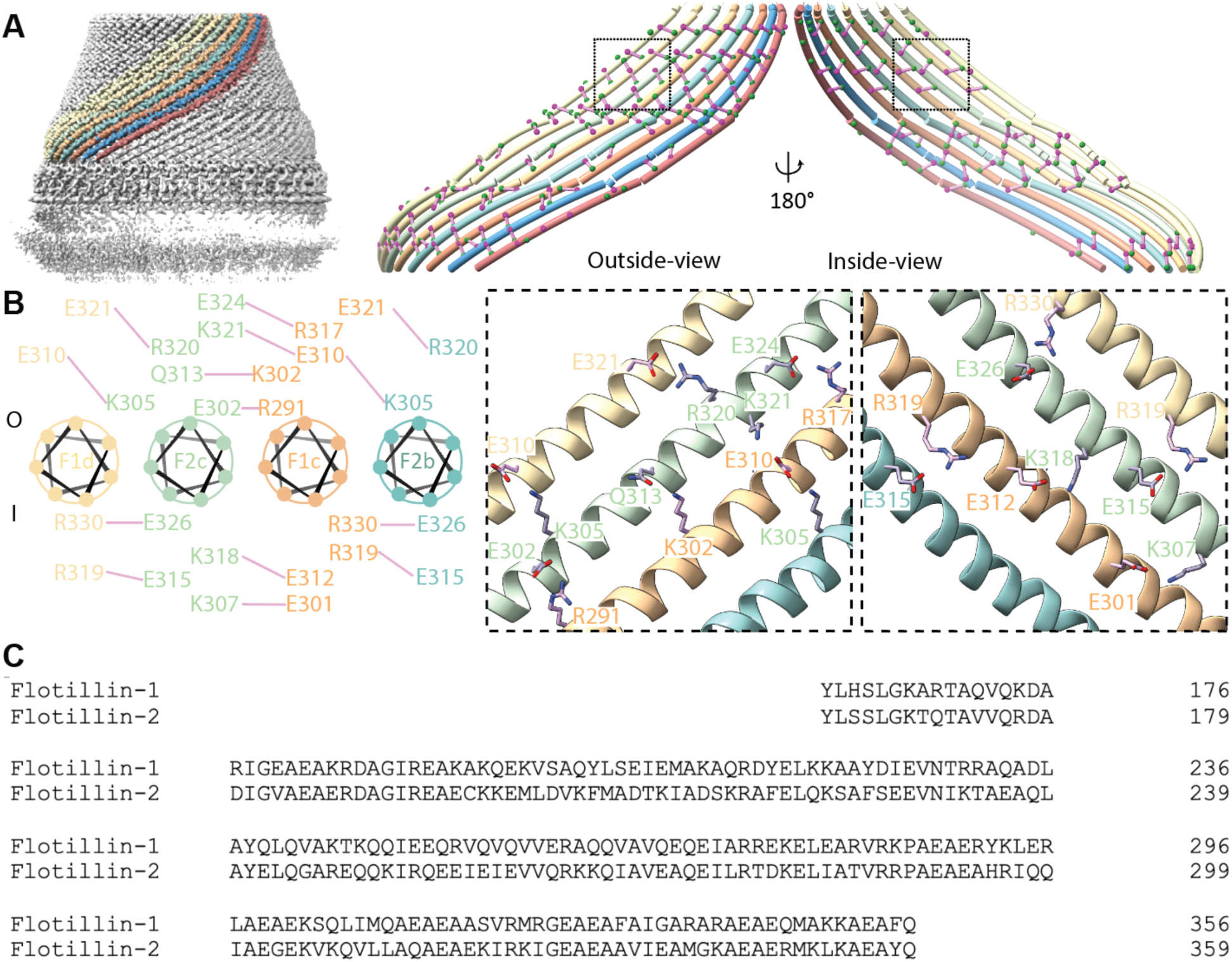
Structure and Organization of Wall Regions in the Flotillin Complex. **(A)** Side view of the Flotillin complex with eight Wall helices separately colored on the left. On the right, both outside-view and inside-view of the eight Wall helices highlight charged residues involved in forming ionized hydrogen bonds (colored in green and magenta), with interactions represented by solid purple lines connecting the alpha-carbons of interacting amino acids. **(B)** Wheel plot illustrating amino acids from four neighboring subunits involved in inter-subunit ionized hydrogen bonds. The top part shows residues outside the Flotillin complex (labeled as O), and the bottom part shows residues on the inside (labeled as I). Dashed box regions correspond to the zoomed-in view from (A), displaying detailed interactions between charged amino acids involved in ionized hydrogen bonding. **(C)** Sequence alignment of Flotillin’s wall helices.

**Fig. S6.**
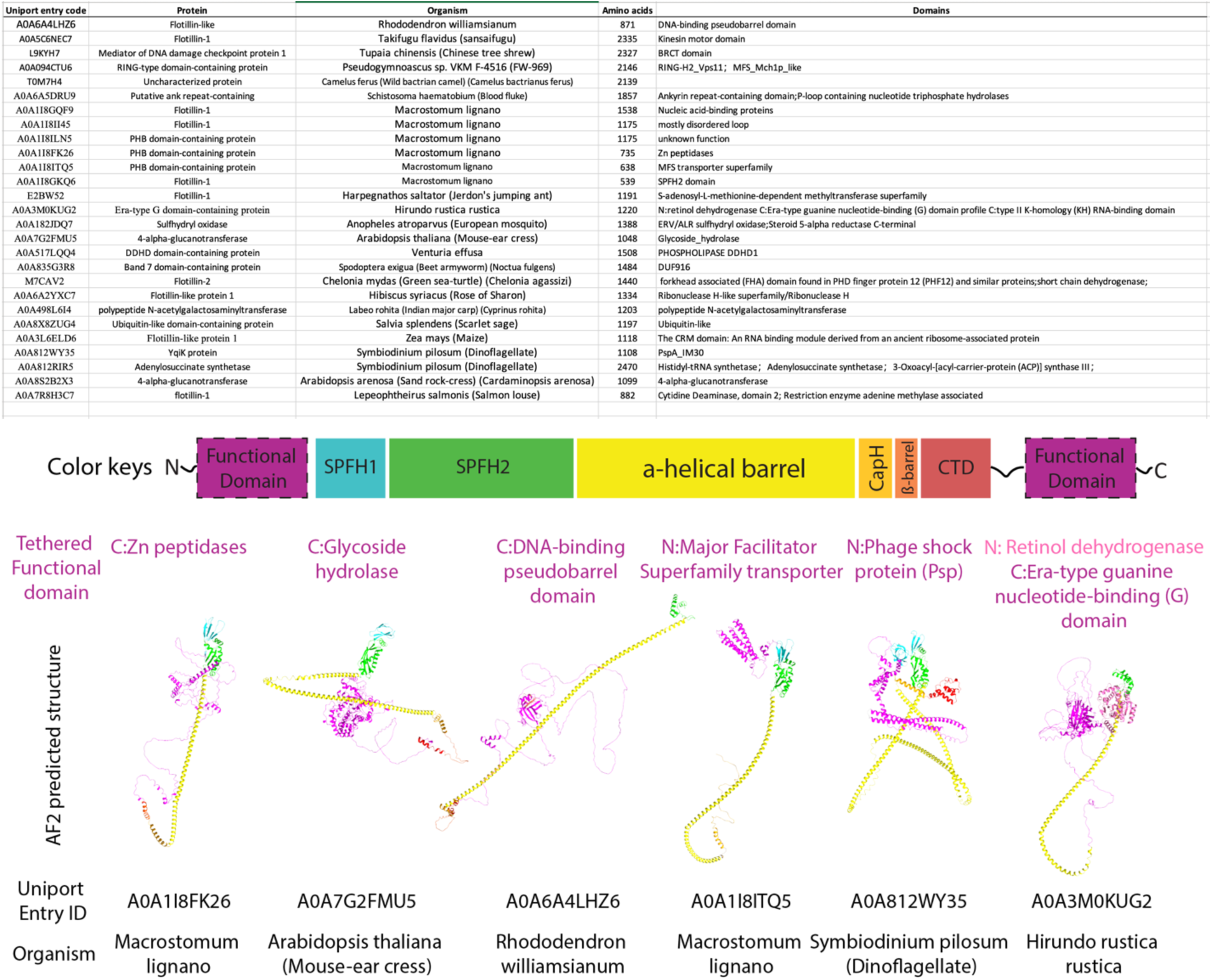
Tethered Functional Domains in Flotillin-Like Proteins. Proteins with conserved Flotillin-like structures and functional domains tethered to their N- or C-termini, as identified in the UniProt database. Each entry includes the UniProt code, protein name, organism, total amino acids, and domain names. The figure illustrates representative Flotillin-like proteins with tethered functional domains; these are colored based on their location at the N- or C-terminus (shown in pink) and labeled accordingly. Below each predicted AlphaFold2 structure, the corresponding UniProt entry ID and organism name are displayed.

**Table S1.**
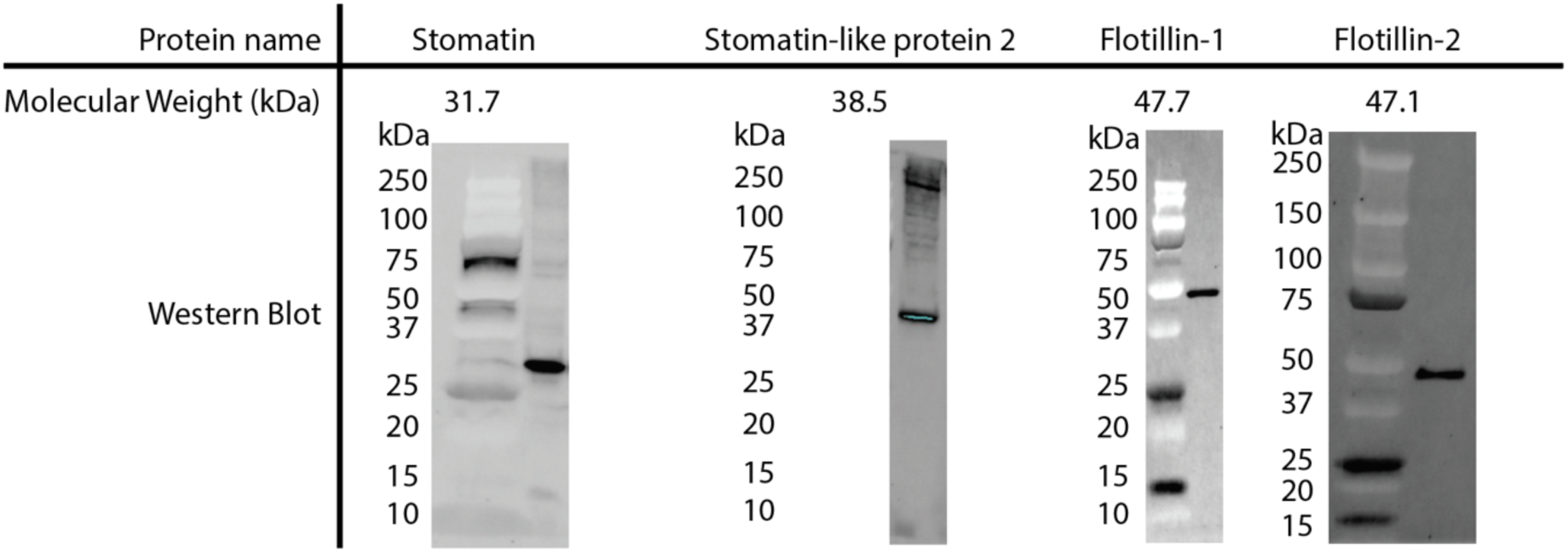
Western Blot Analysis of Stomatin, Stomatin-like Protein 2, Flotillin-1 and Flotillin-2 in HEK293 GnTl^−^ Cells.

**Table S2.**
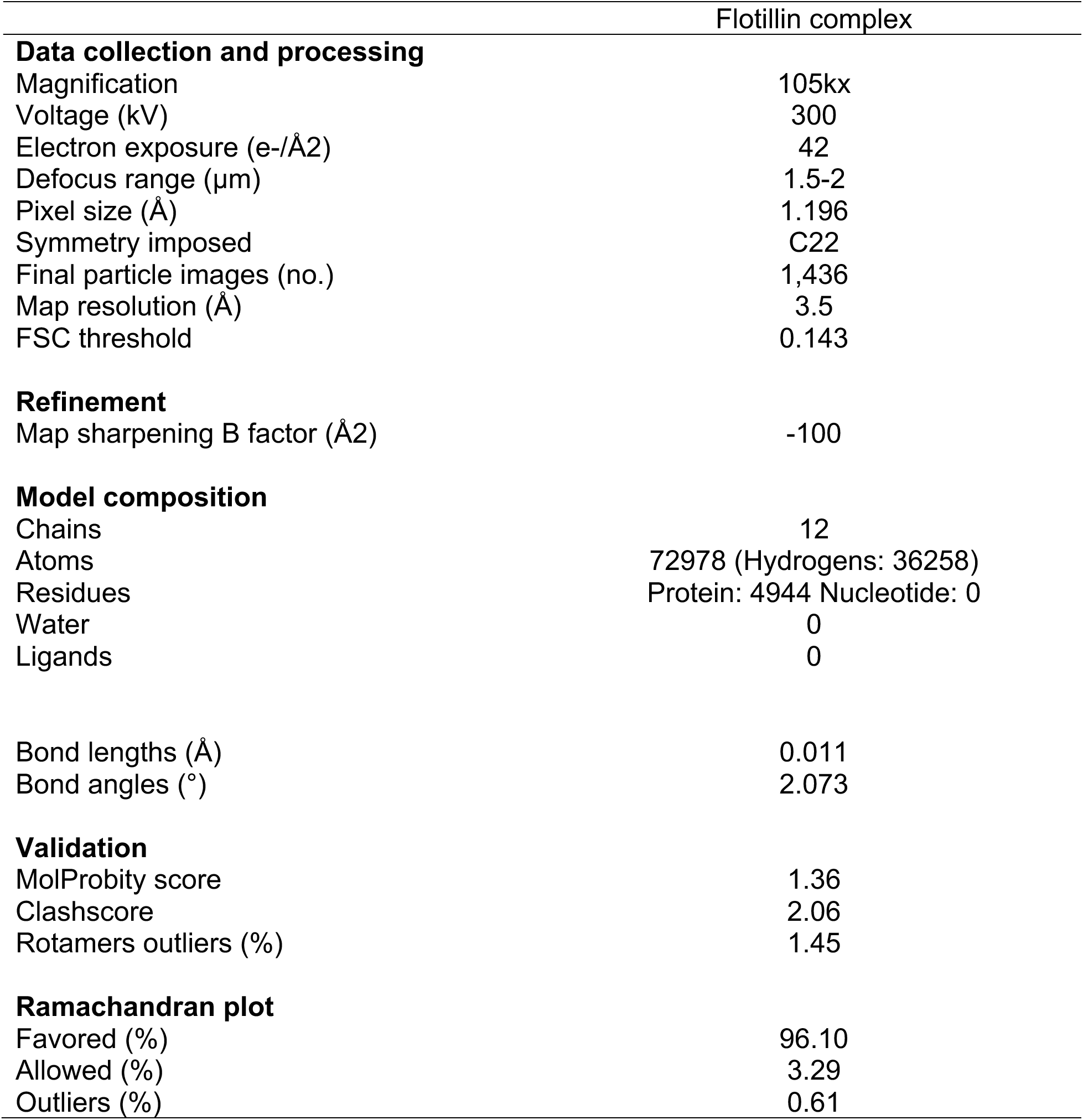
Cryo-EM Data Collection, Refinement and Validation Statistics.

|  | Flotillin complex |
| --- | --- |
| **Data collection and processing** |  |
| Magnification | 105kx |
| Voltage (kV) | 300 |
| Electron exposure (e-/Å<sup>2</sup>) | 42 |
| Defocus range (µm) | 1.5-2 |
| Pixel size (Å) | 1.196 |
| Symmetry imposed | C22 |
| Final particle images (no.) | 1,436 |
| Map resolution (Å) | 3.5 |
| FSC threshold | 0.143 |
| **Refinement** |  |
| Map sharpening B factor (Å<sup>2</sup>) | -100 |
| **Model composition** |  |
| Chains | 12 |
| Atoms | 72978 (Hydrogens: 36258) |
| Residues | Protein: 4944 Nucleotide: 0 |
| Water | 0 |
| Ligands | 0 |
|  |  |
| Bond lengths (Å) | 0.011 |
| Bond angles (°) | 2.073 |
| **Validation** |  |
| MolProbity score | 1.36 |
| Clashscore | 2.06 |
| Rotamers outliers (%) | 1.45 |
|  |  |
| **Ramachandran plot** |  |
| Favored (%) | 96.10 |
| Allowed (%) | 3.29 |
| Outliers (%) | 0.61 |

